# Lipid nanoparticle delivery of siRNA to dorsal root ganglion neurons to treat pain

**DOI:** 10.1101/2025.01.23.633455

**Authors:** Xiangfei Han, Veselina Petrova, Yudong Song, Yu-Ting Cheng, Xingya Jiang, Hui Zhou, Caleb Hu, Dean Shuailin Chen, Hyo Jeong Yong, Hyoung Woo Kim, Biyao Zhang, Omer Barkai, Aakanksha Jain, William Renthal, Philipp Lirk, Clifford J. Woolf, Jinjun Shi

## Abstract

Sensory neurons within the dorsal root ganglion (DRG) are the primary trigger of pain, relaying activity about noxious stimuli from the periphery to the central nervous system; however, targeting DRG neurons for pain management has remained a clinical challenge. Here, we demonstrate the use of lipid nanoparticles (LNPs) for effective intrathecal delivery of small interfering RNA (siRNA) to DRG neurons, achieving potent silencing of the transient receptor potential vanilloid 1 (TRPV1) ion channel that is predominantly expressed in nociceptor sensory neurons. This leads to a reversible interruption of heat-, capsaicin-, and inflammation-induced nociceptive conduction, as observed by behavioral outputs. Our work provides a proof-of-concept for intrathecal siRNA therapy as a novel and selective analgesic modality.

## Introduction

Inadequately treated acute and chronic pain present significant challenges to healthcare systems^1^. Traditional opioid-based and multimodal pain management is limited in many patients by a lack of efficacy, dose-dependent side effects, and public health concerns such as diversion and abuse^2^. Non-pharmacologic adjuvant therapies may improve some aspects of patients’ quality of life, but their evidence-base is weak^3^. Invasive pain therapies, such as intrathecal pumps and spinal cord stimulators, are reserved for very severe and refractory pain states, and are associated with both high cost and the potential side effects^4^. Targeted precision approaches to pain management are few and are currently limited to various monoclonal antibodies against calcitonin gene-related peptide (CGRP)^5^. Other therapeutic targets being investigated include the TRPV1 channel, sodium channels, and nerve growth factor (NGF)^6^, but systemic (e.g., subcutaneous) administration increases their adverse effect profile. These limitations highlight the urgent need for novel, safer and more effective nonopioid therapies for pain management.

Modulating the activity of primary sensory neurons, which transmit both internal and external environmental signals (e.g., thermal, chemical, and mechanical stimuli) to the central nervous system^7, 8^, is an important strategy for pain therapy. The cell bodies of the primary sensory neurons are located in the DRG, providing a unique target at the interface between the peripheral and central nervous system^9^. DRG neurons are pseudounipolar in morphology and constitute a highly heterogeneous population of cells with different cell body diameters, degrees of myelination, receptor expression profiles, and functional characteristics^10-12^. Certain types of primary sensory neurons are known to preferentially detect noxious stimuli (nociceptors) and express unique molecular features compared to neurons that detect innocuous stimuli^10, 13^. For example, sensory neurons expressing the transient receptor potential ion channel, TRPV1 are sensitive to noxious heat (>43 °C), acidic pH, and vanilloid compounds such as capsaicin (the active ingredient in chili peppers)^14, 15^. In preclinical models, TRPV1 genetic ablation results in a marked reduction in noxious heat processing and inflammatory pain induced in the periphery, and therefore represents a potentially favorable target for treatment of these particular pain conditions^14, 16, 17, 18^. However, we currently do not have an effective and versatile strategy for reducing the expression of molecular features in nociceptors (e.g., ion channels like TRPV1) specifically and reversibly and in a manner that avoids systemic side effects.

RNA interference (RNAi) technology, with high specificity and potency in silencing individual genes of interest, has shown immense potential as a novel class of medicine, as exemplified by the clinical approval of multiple siRNA therapeutics^19-21^. However, the success of siRNA therapy has so far been limited to hepatic targeting with the use of LNPs^22^ or N-acetylgalactosamine conjugation^23, 24^. The therapeutic potential of siRNA delivery to DRG neurons for pain treatment has remained largely unexplored^25^. In this study, we thoroughly investigated the use of LNPs, which are closely similar to the formulation used in the FDA-approved Onpattro, for intrathecal siRNA delivery *in vivo*, and revealed key LNP’s physicochemical features required for efficient targeting of DRG neurons. We then evaluated the siRNA LNPs strategy using TRPV1 as a model therapeutic target in mouse models of acute pain, showing efficient and reversible TRPV1 silencing and a corresponding reduction in nociceptive responses to heat-, capsaicin-, and inflammation-induced pain. These findings suggest that intrathecal siRNA LNP delivery to DRG neurons could serve as a robust platform for screening and validating potential nociceptor-selective targets, as well as for developing non-addictive therapeutics for post-traumatic pain or chronic pain induced by diabetes, nerve injury, inflammation, chemotherapy, or genetic mutations.

## Results

To fundamentally explore the intrathecal delivery of siRNA to DRG neurons, we developed a panel of LNPs with varying lipid-PEG compositions, particle sizes, and surface charges (Fig. 1a). All these LNPs comprised four lipid components: ionizable lipid, helper lipid, cholesterol, and lipid-PEG. D-Lin-MC3-DMA (MC3), which is utilized in the clinically approved siRNA LNPs (Onpattro), was chosen as the ionizable lipid^26^, and 1,2-dioleoyl-sn-glycero-3-phosphoethanolamine (DOPE) as the helper lipid. Firefly luciferase siRNA was encapsulated into the LNPs as a model siRNA. Lipid-PEGs have been reported to improve the stability of LNPs and their dissociation kinetics from the LNPs surface affects the protein corona, pharmacokinetics, different cell uptake preference and gene silencing by siRNA LNPs^27-30^. Here, we formulated siRNA LNPs with three commonly used lipid-PEGs with different link groups and tail lengths: 1,2-distearoyl-sn-glycero-3-phosphoethanolamine-n-methoxypolyethylene glycol (DSPE-PEG), 1,2-distearoyl-rac-glycero-3-methoxypolyethylene glycol (DSG-PEG), and 1,2-dimyristoyl-rac-glycero-3-methoxypolyethylene glycol (DMG-PEG), with the same PEG molecular weight of 2,000 and methoxy terminal group. Similarly, particle size and surface charge of LNPs have also shown dramatic impact on their tissue penetration, pharmacokinetics, biodistribution, and siRNA gene silencing potency^31-35^. We thus prepared LNPs with different average sizes (55 nm, 92 nm, and 153 nm) by changing the DMG-PEG ratio of the LNP formulations (Supplementary Fig. 1), as well as different surface charges (+15.2 mV, −23.9 mV, and −2.1 mV) by using DMG-PEG with various terminal groups (DMG-PEG-NH2, DMG-PEG-COOH, and DMG-PEG-OCH3) (Supplementary Table 1).

**Fig. 1:**
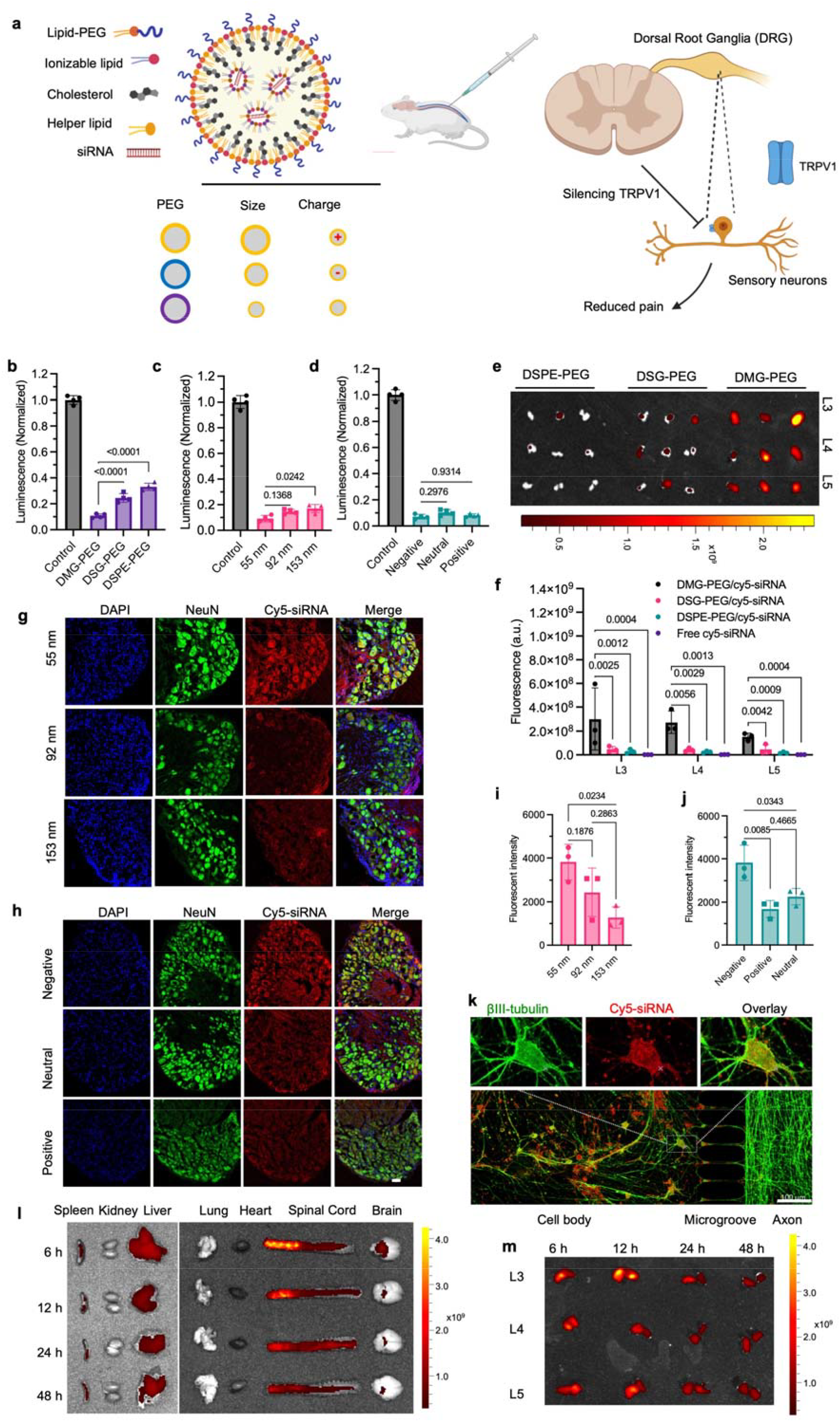
LNPs enable efficient intrathecal siRNA delivery to DRG. **a**, Schematic illustration of LNPs formulated for DRG targeting. siRNA-loaded LNPs were prepared with ionizable lipid (MC3), helper lipid (DOPE), cholesterol, and lipid-PEG. A panel of 9 different LNP formulations was prepared, by varying lipid-PEG composition, particle size, and surface charge. Lead siRNA LNPs were selected for intrathecal injection between lumbar L4 and L5 to investigate the analgesic effect in pain models. **b-d**, Effect of lipid-PEG composition (**b**), particle size (**c**), and surface charge (**d**) on luciferase silencing in 175-luc cells *in vitro* (N=4, mean ± SD, One-way ANOVA with Tukey’s post hoc test). **e**, Effect of lipid-PEG on cy5-siRNA LNP accumulation in the DRG. C57BL/6J mice were administered with different siRNA-LNP formulations by intrathecal injection, followed by L3-L5 DRGs imaging 24 hrs post-administration. **f**, Quantitative analysis of cy5 fluorescence intensity in DRG neurons from **e. g**,**h**, Effect of LNP size (**g**) and surface charge (**h**) on cy5-siRNA accumulation in DRG neurons. Mice were administered with cy5-siRNA LNPs via intrathecal injection, followed by imaging of L3-L5 DRG tissue section 24 hrs post-administration. **i**,**j**, Quantitative analysis of cy5 fluorescence intensity in DRG neurons from **f** and **g**, respectively (N=3, mean ± SD, One-way ANOVA with Tukey’s post hoc test). **k**, βIII-tubulin immunostaining and confocal imaging of DRG neurons and axons treated with cy5-siRNA LNPs in the axon compartment. Scale bar: 100 µm. **l**,**m**, *In vivo* biodistribution of cy5-siRNA LNPs in major organs (**l**) and lumbar L3-L5 DRGs (**m**) at different time points following intrathecal injection. The unit for IVIS fluorescence intensity in **e, l**, and **m** is (p/sec/cm^2^/sr)/(μW/cm^2^).

We first measured *in vitro* the luciferase silencing efficiency of these LNPs in a luciferase-expressing RIL-175 (175-luc) mouse cell line using the Steady-Glo Luciferase Assay. Among these formulations, LNPs composed with DMG-PEG achieved the strongest luciferase silencing (>90%) compared to those with DSG-PEG (~80%) and DSPE-PEG (~70%) at an siRNA concentration of 20 nM (Fig. 1b). Additionally, the smallest particle size (55 nm) yielded the highest silencing efficiency over 90%, while 92 nm and 153 nm LNPs enabled ~80% silencing (Fig. 1c). We also tested the effect of LNP surface charge on luciferase silencing *in vitro*. Fig. 1d shows that there was no significant difference in luciferase silencing among these groups. We then studied the cytotoxicity of these LNPs in 175-luc cells with an AlamarBlue assay, and no significant cytotoxicity was observed following treatment (Supplementary Fig. 2). We further investigated cellular uptake and endosomal escape by culturing 175-luc cells with cy5-siRNA-loaded LNPs. The cellular fluorescent intensity was measured using both flow cytometry and confocal microscopy (Supplementary Figs. 3 and 4). LNPs with DMG-PEG and smaller particle size enabled higher uptake than other groups. For the surface charge, there was no significant difference between three different charged LNPs. We then quantified the endosomal escape by calculating the correlation efficiency of endosome with cy5-siRNA, and we found no significant difference among the different formulations (Supplementary Fig. 5). These results indicate that the luciferase silencing differences among these LNPs may be attributed mainly to cellular uptake rather than endosome escape.

We also explored the targeting efficiency of these LNPs into DRG neurons *in vivo*. Mice were administered cy5-siRNA-loaded LNPs via intrathecal injection and lumbar L3-L5 DRGs were collected for *in vivo* imaging 24 hrs post-administration. LNPs coated with DMG-PEG enabled the highest accumulation of cy5-siRNA in the lumbar DRGs. In contrast, DSG-PEG LNPs and DSPE-PEG LNPs showed significantly less DRG accumulation (Fig. 1e,f). Notably, compared with free cy5-siRNA, DMG-PEG LNPs, DSG-PEG LNPs, and DSPE-PEG LNPs enabled 204.8-, 32.5- and 19.5-fold higher accumulation of cy5-siRNA in mouse L3 DRGs, respectively (Fig. 1f, Supplementary Fig. 6). DMG-PEG LNPs also induced 107.1-fold and 108.5-fold higher accumulation of siRNA in L4 and L5 DRGs, than free cy5-siRNA. These results suggest that LNPs could dramatically improve siRNA delivery to DRGs and that lipid-PEG significantly impacts the distribution of LNPs.

We then conducted immunofluorescent staining on cryosectioned DRGs to investigate the effect of LNP size and charge on DRG neuron targeting. After intrathecal administration, cryosectioned DRGs were labeled with an anti-NeuN antibody (a neuronal marker) and imaged using confocal microscopy. LNPs with a 55 nm size and a negative surface charge achieved the highest DRG neuron accumulation (Figs. 1g-j). The 55 nm LNPs showed ~1.5 and ~3-fold greater fluorescent intensity increases than the 92 nm and 153 nm LNPs, respectively. In addition, anionic LNPs showed ~2-fold fluorescent greater intensity increases than positive and neutral LNPs. Notably, there is no significant difference between positive and neutral LNPs. Smaller particle size and negative charge enable high DRG neuron uptake of cy5-siRNA LNPs, which might be related to better tissue diffusion^35, 36^ To further confirm the LNP’s size and charge effects on DRG targeting, we collected L3-L5 DRGs from healthy mice and incubated them with the cy5-siRNA LNPs in the culture medium. Then we imaged the ex vivo DRGs using an IVIS imaging system. We found that LNPs with small particle size (55 nm) and negative charge showed the highest cy5-siRNA accumulation (Supplementary Fig. 7), which is fully consistent with the *in vivo* targeting efficiency results. Those LNPs with a size of 55 nm and a negative DMG-PEG-COOH surface coating (referred to as 55NM LNPs) showed the most efficient DRG neuron delivery after intrathecal injection. To have a better understanding of LNP transportation to DRG neuron bodies, we investigated the axon transport of the 55NM LNPs *in vitro*. We cultured mouse primary DRG neurons in a Xona microfluidic chamber and allowed axons to grow from the cell body compartment across the microchannel and reach the axonal compartment^37^. Then cy5-siRNA-loaded 55NM LNPs were added to the axon compartment and the whole chamber was imaged by confocal microscopy 24 h post-treatment. Fig. 1k showed that 55NM LNPs were actively transported along axons in a retrograde manner and reached the cell bodies of DRG neurons. This indicates that 55NM LNPs could be transported via axons to neuron bodies, in addition to direct diffusion into DRGs. We further assessed the distribution of cy5-siRNA in major organs at various time points post-intrathecal injection. cy5-siRNA mainly accumulated in the spinal cord and DRGs, with peak fluorescence intensity in lumbar DRGs observed ~6-12 hrs post-administration, followed by a gradual decrease over time (Fig. 1l,m).

Efficient gene silencing in DRG neurons could be a promising approach to treat peripherally driven pain. Here, we chose TRPV1 as a proof-of-concept and clinically relevant target to investigate the efficacy of the TRPV1 siRNA-loaded 55NM LNPs (55NM/TRPV1) *in vitro* and *in vivo*. Scrambled siRNA-loaded 55NM LNPs (55NM/SCR) and PBS were used as controls. TRPV1 mRNA levels of 175-luc cells and DRGs were measured by quantitative reverse transcription polymerase chain reaction (RT-qPCR) *in vitro* and after intrathecal injection, respectively. TRPV1 mRNA expression was reduced to approximately 10% of baseline levels 2 days post-treatment, and recovered partially by 1 week (~56% expression) and fully at 4 weeks (Fig. 2a,b). This short-term silencing effect would be perfectly suited to treat acute pain (e.g., perioperatively), avoiding prolonged alterations in pain sensation.

**Fig. 2:**
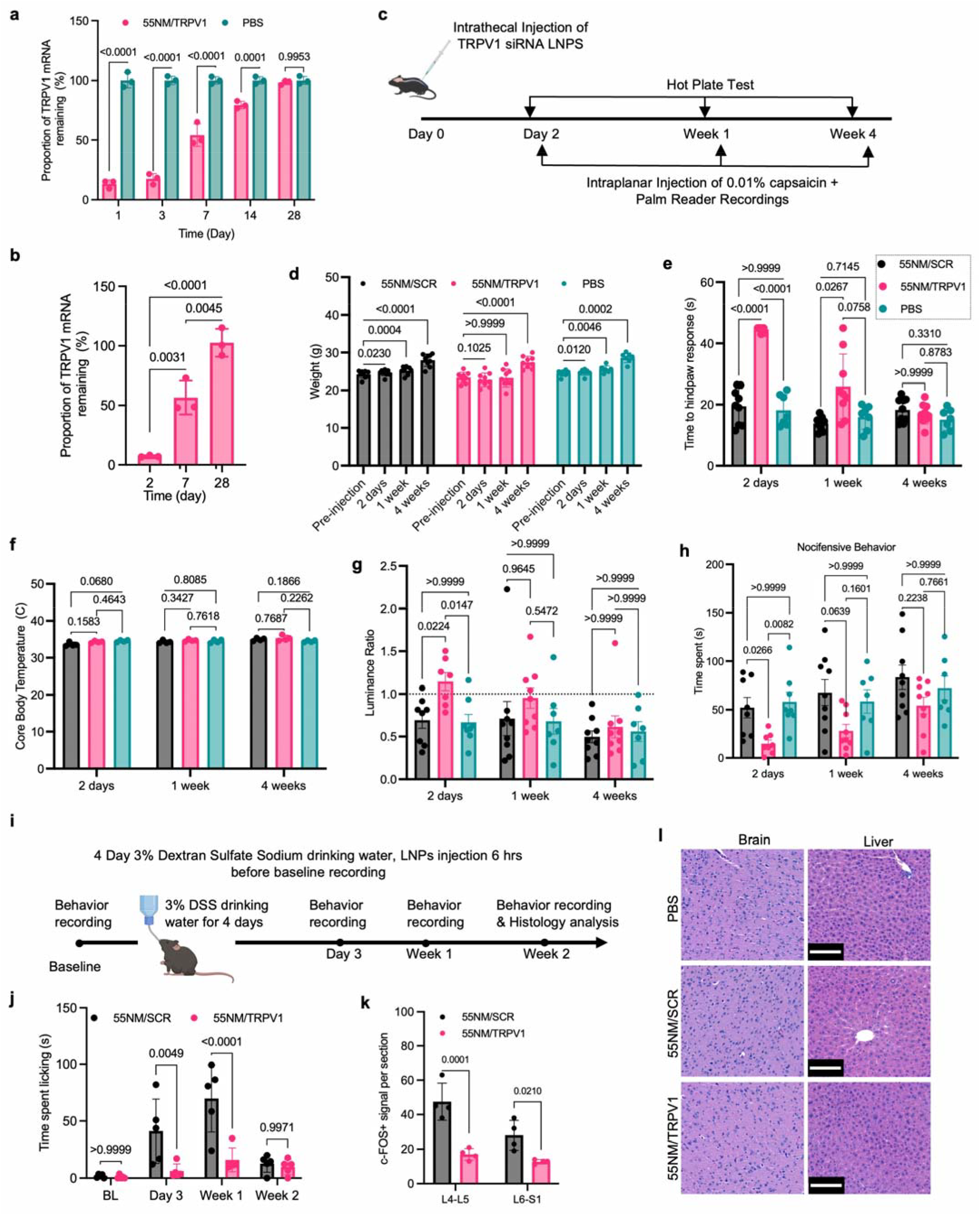
TRPV1 siRNA LNPs mediate robust gene silencing and reduce noxious heat and capsaicin-induced acute pain. **a**, TRPV1 mRNA expression in 175-luc cells after treatment with TRPV1 siRNA-loaded 55NM LNPs for 24 h. (n=3, mean ± SD). Total RNA was extracted at different time points and TRPV1 mRNA levels normalized with GAPDH mRNA. **b**, TRPV 1 mRNA expression in the DRG of mice following intrathecal administration of TRPV1 siRNA LNPs (n=3, mean ± SD). **c**, Timeline of animal experiments. **d**, Changes in body weight of mice before and after the treatments. **e**, Heat withdrawal latency of treated mice with TRPV1 siRNA LNPs using hot plate test. Mice injected with 55NM/TRPV1 LNPs (n=9) exhibited significantly prolonged response times to noxious heat compared to PBS-treated (n=7) or 55NM/SCR-treated mice (n=9) 2 days post-delivery (Two-way ANOVA with Bonferroni post hoc test). This effect was partially reduced at 1 week and absent by 4 weeks post-delivery. **f**, No significant changes in core body temperature were observed in mice after receiving 55NM/SCR LNPs, 55NM/TRPV1 LNPs or PBS (Two-way ANOVA with Bonferroni post hoc test). **g**, Luminance ratio of the left-to-right hind paw in mice treated with TRPV1 siRNA LNPs. Luminance ratio was significantly increased in mice treated with 55NM/TRPV1 LNPs (n=8, ratio=1.1) compared to those receiving 55NM/SCR LNPs (n=7, ratio=0.69) or PBS (n=8, ratio=0.67), 2 days post-delivery. This effect diminished at 1 and 4 weeks post-delivery (Two-way ANOVA with multiple comparisons, Bonferroni post hoc test). **h**, Nocifensive behavior of mice treated with TRPV1 siRNA LNPs. Mice administrated with 55NM/TRPV1 LNPs showed a significantly reduced duration of nocifensive behavior (n=8) compared to 55NM/SCR (n=8) or PBS controls (n=7, 58 s), 2 days post-delivery (Two-way ANOVA with multiple comparisons, Bonferonni test). This effect diminished at 1- and 4-weeks post-administration. **i**, Experiment timeline of DSS-induced inflammatory pain and spontaneous behavior recording at 3 days, 1 W and 2 W post LNP treatment. **j**, Temporal profile of time spent licking anal region in mice treated with TRPV1 siRNA LNPs and scrambled LNPs. Mice administered with 55NM/TRPV1 LNPs exhibited significantly reduced anal licking compared to 55NM/SCR-treated mice (n=5) on Day 3 and 1W post-treatment (Two-way ANOVA with Šídák’s post hoc test). **k**, Quantification of cFOS^+^ signal in spinal cord at segments L4-L5 and L6-S1 in two groups on Day 14 post-treatment (n=4 mice per group, Two-way ANOVA with Šídák’s post hoc test). **l**, Hematoxylin and eosin (H&E) staining of brain and liver post-administration with different formulations showed no significant histopathological changes. Scale bar = 100 µm.

Motivated by the robust *in vitro* and *in vivo* gene silencing results, we explored the effect of TRPV1 silencing on three different behavioral paradigms: the response to noxious heat-, capsaicin-, and prolonged inflammation-induced pain. In the first set of experiments, 10 µl TRPV1 siRNA LNPs (55NM/TRPV1) or scrambled siRNA LNPs (55NM/SCR) were administered intrathecally to mice at a dose of 0.15 mg/kg siRNA and noxious heat responses assessed at 2 days, 1 week, and 4 weeks post-administration (Fig. 2c). 55NM/TRPV1 treatment did not adversely affect the overall health or weight of the animals (Fig. 2d). In the hotplate test^38^, the paw is heated by contact with a hot metal surface and the time to paw licking or withdrawal is measured. TRPV1 knockdown via siRNA LNPs delivery practically abolished the responses to noxious heat (52 °C) 2 days post-administration, as the mice did not respond to the noxious heat within the 45 second limit (Fig. 2e). This indicates robust deficits in thermally evoked pain-related behavior. This analgesic effect began to diminish by 1 week and was fully resolved by 4 weeks post-administration, aligning with the restoration of TRPV1 mRNA levels (Fig. 2b). Importantly, TRPV1 knockdown by siRNA LNPs did not result in changes in core body temperature (Fig. 2f), unlike systemic TRPV1 antagonists which are under clinical investigation but have major side effects such as hyperthermia^39^.

Next, we evaluated capsaicin-evoked pain behavior by applying 0.01% capsaicin to the left hind paw after intrathecal treatment of 55NM/TRPV1 (Fig. 2c). Mice were placed in a black box chamber where they could explore freely and near-infrared frustrated total internal reflection was used to detect the degree, force and timing of the paw surface contact as well as body pose measures of naturalistic mouse behavior^40^. In naïve mice, the paw luminance ratio is near 1 because the mice place equal force on both hind paws^40^. When capsaicin is injected, the paw luminance ratio falls below 1, as mice avoid putting force on the sensitized paw. In 55NM/TRPV1-treated mice, the paw luminance ratio was significantly higher on day 2 (ratio = 1.1) compared to that of 55NM/SCR-treated mice (ratio = 0.69) and PBS-treated mice (ratio = 0.67) (Fig. 2g), indicating absence of mechanical pain-hypersensitivity response to the capsaicin-mediated activation of TRPV1-expressing neurons. In addition, 55NM/SCR- and PBS-treated mice exhibited nocifensive behaviors (licking, biting, and flinching) for an average of 52 and 58 sec, respectively, within a 3-min observation period (Fig. 2h). In contrast, 55NM/TRPV1-treated mice displayed significantly reduced nocifensive behaviors (~15 sec) 2 days post-delivery, which is comparable to transgenic shRNA expression^41^. This effect was reduced at 1 week and diminished 4 weeks after a single injection, which signifies a relatively short-acting analgesic effect.

We further evaluated the effect of 55NM/TRPV1 treatment on spontaneous pain-like behaviors at 3 days, 1 week, and 2 weeks post-administration in mice that had been subjected to 4-days of 3% dextran sulfate sodium (DSS) in their drinking water. This treatment induced gut inflammation and pain (Fig. 2i), which leads to diarrhea and triggers the mice to lick their anal port^42^. In 55NM/SCR-treated mice, the spontaneous licking behaviors peaked at day 7 (3 days after the DSS was removed from the drinking water) and gradually subsided over 2 weeks. In mice treated with 55NM/TRPV1, nocifensive behaviors showed a 7-fold and 4-fold reduction of time spent on licking at day 3 (~6 sec vs. ~41 sec) and day 7 (~16 sec vs. ~70 sec), respectively (Fig. 2j, Supplementary Fig. 8). To further verify the analgesic effect of the LNP treatment on silencing the neuronal activation induced by the DSS induced peripheral colonic inflammation, we performed cFOS staining, a biomarker for neuronal activity, in the cryosectioned spinal cord segments at lumbar 4-5 (L4-L5) and at lumbar 6 and sacral 1 (L6-S1) to assess treatment efficacy. We found that there was significantly reduced cFOS positive signals in the spinal cord of the group with 55NM/TRPV1 treatment compared with the 55NM/SCR-treated group (Fig. 2k). This approach has enabled us to explore the functional consequences of short-term gene silencing without compromising long-term gene function, which is critical for maintaining health and development, as well as the capacity to detect and react to potentially damaging stimuli. Notably, 55NM/TRPV1 produced behavioral phenotypes similar to TRPV1 knockout mice^41, 43^ and TRPV1 antagonist treatment^44-46^, but with reduced side effects such as hyperthermia^47^, locomotion deficits, and aberrant inflammation, and with a predictable reversibility.

To assess the *in vivo* safety of the siRNA LNPs we conducted a panel of serum metabolic studies, including alkaline phosphatase (ALP), alanine aminotransferase (ALT), and aspartate aminotransferase (AST). We did not find observable blood toxicity or hepatic function damage (Supplementary Fig. 9). We also performed histological analysis on paraffin-embedded sections of major organs (brain, liver, heart, spleen, lungs, and kidneys) using Hematoxylin and Eosin (H&E) staining. There is no apparent organ damage or inflammatory lesions following intrathecal injection of 55NM/TRPV1 in comparison to the PBS group (Fig. 2i, Supplementary Fig. 10). These findings highlight the potential of siRNA LNP delivery for alleviating acute and inflammatory pain safely and efficiently through TRPV1 silencing in the DRG.

Some limited efforts were previously made with intrathecal siRNA delivery to DRGs for studying or evaluating pain pathways^48-52^. These studies adopted a daily intrathecal pump injection^50^ or cationic lipid/polymer transfection agents^48, 49, 51, 52^, which are not feasible for translation given their low efficiency or toxicity concerns. The intrathecal delivery of translatable siRNA LNPs to DRG neurons has not been explored before. Our work suggests a promising therapeutic potential of intrathecal siRNA LNPs as analgesics. With recent advancements in our understanding of the pathophysiology of pain and the identification of underlying molecular mechanisms, one can envision a precision targeting of these mechanisms at a molecular level as an effective way to reduce pain. A rise in NGF levels after peripheral nerve injury has been associated with ion channel sensitization and pain. Therefore, receptors such as TrkA, the NGF receptor could be an attractive target for the transient suppression of NGF’s pain enhancing actions after injury. In addition, suppression of Nav1.7 and Nav1.8 could also result in acute and chronic pain alleviation. Targeting such channels has to be done in a controlled and reversible manner to ensure that the protective function of pain is preserved. Other channels such as TRPA1 and TRPV4 have been implicated in diabetic neuropathy pain and could also represent future potential targets^53-55^. Immune receptors such as IL-10, TLR4 and IL-1 represent a set of promising targets for treating chemotherapy-induced peripheral pain. However, further investigation is needed to confirm that no side effects will be observed due to the widespread expression of these immune receptors on many different cell types and tissues^56-59^. Recent advances in single-cell RNA-sequencing have significantly expanded the list of channels and receptors that are preferentially expressed in nociceptors and could serve as safe and effective analgesic targets^10, 12, 13, 60- 62^. The ease of designing siRNAs to any possible target and encapsulating them in LNPs of specific size and charge, as well as the relatively rapid onset of gene silencing and the wide range of pain behaviors that can be examined, make the siRNA LNP platform highly suitable for *in vivo* screening and advancing our understanding of pain molecular mechanisms and the discovery of novel analgesics.

In summary, we revealed a dramatic improvement in intrathecal siRNA delivery to DRGs neurons by using LNPs, as well as the impact of the physiochemical properties of the LNPs on DRG neuron targeting and gene silencing efficiency. Our 55NM LNPs demonstrated the ability to successfully deliver siRNAs to lumbar DRGs *in vivo* and selectively silence specific genes to elicit excellent analgesic effects in different mouse models. While our findings highlight the therapeutic potential of siRNA LNPs for pain treatment, further refinement in LNP formulation and siRNA design/modification may be required to ensure safe and effective clinical applications. A comprehensive evaluation of the efficacy profile of this approach on different acute and chronic pain models such as diabetes, chemotherapy, and nerve injury-induced neuropathic pain is now required. This project has advanced the understanding of DRG-targeted siRNA delivery and establishes a foundation for exploring broader applications for the treatment of neuropathic and inflammatory pain disorders. The siRNA LNP strategy may provide an opportunity for the development of safe, effective, and non-addictive novel treatments of pain.

## Materials

D-Lin-MC3-DMA (MC3) was purchased from BroadPharm. 1,2-dioleoyl-sn-glycero-3-phosphoethanolamine (DOPE), cholesterol, and 1,2-dimyristoyl-rac-glycero-3-methoxypolyethylene glycol-2000 (DMG-PEG-OCH3) were purchased from Avanti Polar Lipids. DMG-PEG-NH2 and DMG-PEG-COOH with PEG MW of 2,000 were purchased from Nanosoft Polymers. Luciferase siRNA, cy5-luciferase siRNA (cy5-siRNA), and TRPV1 siRNA were purchased from Horizon Discovery.

### Formulation and characterization of siRNA LNPs

MC3, DOPE, cholesterol, and lipid-PEG were used to prepare siRNA LNPs. The details of lipid composition for the formulation of different siRNA LNPs are shown in Supplementary Table 1. The siRNA cargoes were dissolved in 10 □ mM citric buffer (pH 3), formulated with the lipids in ethanol using pipetting or iNano E microfluidic mixing, and dialyzed against 1× □ PBS in 10 kDa Puri-A-Lyzer™ Midi Dialysis Kit (Sigma) at 4 □°C for overnight. siRNA LNPs were concentrated after dialysis using a MWCO 10 □ kDa Amicon Ultra-4 mL centrifugal filter unit for *in vivo* studies. LNPs’ size distribution and surface potential were determined using ZETAPals analyzer (Brookhaven Instruments, Holtsville, NY) at 25□°C.

### *In vitro* luciferase silencing

Luciferase-expressing RIL-175 (175-luc) cells were seeded in 96 well plates and treated with different siRNA LNPs (20 nM) for 24 hrs. After incubation, the cells were washed with fresh medium and the expression of firefly luciferase in 175-luc cells was determined using the Steady-Glo luciferase assay kit. The luminescence intensity was measured using a Tecan microplate reader, and the average value of independent measurements was calculated.

### Cellular uptake of cy5-siRNA LNPs

175-luc cells were cultured in Dulbecco’s modified Eagle’s medium (DMEM, cat. #11965175, Thermo Fisher Scientific) supplemented with 10% heat-inactivated fetal bovine serum (FBS, Hyclone), 1× penicillin/streptomycin. Flow cytometry was used to confirm the cellular uptake of the LNPs. Approximately 30,000 cells were seeded per well in 12 well plates and incubated at 37□°C overnight in a 5% CO_2_ atmosphere. Cells were treated with 20 nM cy5-siRNA LNPs and incubated for 24□hrs. Following incubation, cells were washed twice with PBS, trypsinized using Trypsin (0.25%)-phenol red (Gibco, cat. #15050065), harvested and re-suspended in the FACS buffer, followed by flow cytometry analysis (10,000 cells count, BD Biosciences). Data were processed using FlowJo 10.

### *In vitro* cell viability

A cell density of 7,000 per well was plated in white, clear-bottom 96-well plates and allowed to grow to 60-70% confluency overnight at 37 confluency overnight °C. 175-luc cells were treated with various siRNA LNPs at the concentration of 20 nM. All the cells were incubated for an additional 24 hrs and cell viability was measured with the AlamarBlue assay.

### Endosomal escape study

175-luc cells at a density of 60,000 per 8-well (Thermo Fisher Scientific) were incubated overnight for attachment. Cy5-siRNA LNPs were then added to the cells and incubated for 24 □ hrs. After that, media was aspirated, and cells were washed with 1x DPBS and stained with LysoTracker™ Green for 30mins. Then cells were washed with PBS and fixed with 4% paraformaldehyde for 15 □ minutes at room temperature. After fixing, cells were washed 3 times with 1x DPBS, stained with DAPI and mounted using ProLong™ Gold Antifade Mountant (Invitrogen). Images were taken with a Zeiss 710 confocal microscope. Endosome escape was quantified by calculating the correlation coefficients between cy5 and endosome using ImageJ.

### Collection of primary DRG neurons

L3-L5 DRGs were removed from mice and placed in filtered ice-cold Hanks’ Balanced Salt solution (HBSS) until dissociation. The DRGs were placed in 1 mL Neurobasal-A medium containing Collagenase A (Roche, 5mg/ml) and DispaseII (Roche, 1mg/ml) at 37 °C for 70 min. Next, the DRGs were washed twice with neuron culture media (Neurobasal-A medium containing 1% penicillin–streptomycin, 1% GlutaMax supplement, and 2% B-27 supplement). After that, they were mechanically dissociated by successive cycles of up and down aspiration cycles through a pipet to generate cell suspensions, which were then filtered through a 70 µm cell strainer to a new tube. The solution was then centrifuged, replaced with 10% BSA solution, and centrifuged again. DRG neurons were collected from the interphase and went through another centrifuge and neuron culture medium replacement before being plated.

### Axon transportation of siRNA LNPs

A microfluidic device (XC150, Xona Microfluidics) was used to achieve compartmental culture of DRG neurons. It consists of two chambers connected with microfluidic channels. Prior to seeding DRG neurons, the microfluidic device was coated with poly-D-lysine (1 mg mL^−1^) and laminin (30 µg mL^−1^) sequentially. After coating, the microfluidic device was placed in an incubator (37 °C, 5% CO_2_) until ready to seed DRG neurons. A cell suspension of dissociated DRG neurons was prepared to obtain a density of 1 × 10^6^ cells mL^−1^. 10 µL of cell suspension was seeded only in the left chamber of the microfluidic device (cell body side). The microfluidic device was placed in an incubator (37 °C, 5% CO_2_) for 30 min to allow the cells to attach. Then, 150 µL of neuron culture media was added to each of the two chambers, and the microfluidic device was placed into 37 °C, 5% CO_2_incubator again. Glial cell line-derived neurotrophic factor (GDNF) and nerve growth factor (NGF) were added to the cell body compartment at 50 ng/ml and to the axon compartment at 100 ng/ml for 1 day and changed to 10 ng/ml (cell bodies) and 100 ng/ml (axons) and maintained for 5-10 days. During this time, axons extended from the cell body side and grew through the microfluidic channels, reaching the axon side. The microfluidic device remained in the incubator to ensure sufficient axonal growth on the axon side. Then cy5-siRNA LNPs were added to the axon compartment of the microfluidic device and a control neuron culture medium was added to the cell body side. The chamber was cultured in cell incubator overnight. A small volume (50 µL) difference between the cell body side (150 µL) and the axon side (100 µL) was maintained to prevent passive diffusion of the siRNA LNPs across the microfluidic channels. Then, both sides were washed with PBS, fixed using 4% paraformaldehyde for 30 min, rewashed with PBS again, and blocked with a blocking solution (PBS containing 5% BSA, 0.4% Triton-X, 0.05% Tween 20) for 30 min. The DRG neurons were then incubated with the primary antibody anti-βIII tubulin (Sigma cat# T3952, 1:500) overnight in PBS containing 3% BSA, 0.4% Triton-X, 0.05% Tween 20 at 4 °C, washed with PBS, incubated with the secondary antibody (goat anti-rabbit Alexa Fluor™ 546 Invitrogen A-11035, 1:1000 dilution) for 1 h, and washed with PBS again. The DRG neurons were then stored in PBS containing 4% paraformaldehyde solution at 4 °C before confocal fluorescence imaging.

### Animal studies

All animal experiments were conducted in accordance with protocols approved by the Institutional Animal Care and Use Committee (IACUC) at Brigham and Women’s Hospital and Boston Children’s Hospital. 8-week-old C57BL/6J mice were obtained from the Jackson Laboratory (JAX:000664). All animals were housed in a pathogen-free facility with free access to food and water, at a standard 12-hr light/dark cycle. Animals were randomized to treatment groups. Intrathecal injections (5 µL in volume) were performed on anesthetized and shaved animals, using a 30 gauge, sterile, single-use needle that was inserted through the skin over L4-L5 of the spinal column. Intraplantar injection of 0.01% capsaicin solution (Tocris) to the left paw was performed using a 31-gauge, sterile, single-use needle. Inflammatory pain was induced by feeding 3% of DSS (MP Biomedicals 0216011080) into the regular drinking water for 4 days in mice.

### Immunofluorescence staining of DRG and spinal cord tissues

The mouse lumbar DRGs and spinal cords were collected for immunofluorescence imaging. Animals underwent transcardiac perfusion, and harvested tissues were placed in 4% paraformaldehyde overnight. After fixation, whole globes were rinsed in PBS and then incubated in 30% sucrose for 2 □ days at 4 °C before embedding in cryostat compound (Tissue-Tek O.C.T. Compound, code # 4583). The embedded DRGs were snap-frozen in a dry ice bath. DRGs were sectioned at 12 microns and spinal cords were sectioned at 40 microns with a cryostat (Leica 1950) and stored at −20 □ °C. For immunostaining, slides were washed in PBS for 15 mins, followed by 3x 10-min washing with 1% Triton-X. Slides were incubated in blocking buffer for 2 hrs at room temperature (5% BSA, 0.4% Triton-X, 0.05% Tween 20) and then incubated with primary antibody overnight at 4 °C (Rabbit anti-NeuN Abcam cat# 1:500; Rabbit anti-cFOS Cell Signaling Technology cat# 2250 1:200). The next day, slides were washed 3x with PBS and incubated with secondary antibody for 2 hrs at room temperature (Donkey anti-Rabbit Alexa Fluor™ 488, A-21206, 1:1000; Goat anti-Rabbit Alexa Fluor™ 594 Invitrogen A-11012 1:500). Slides were then washed 3x with PBS and mounted with Prolong antifade DAPI medium (Invitrogen, #P36935). Confocal images were taken with a ZEISS LSM 710 confocal microscope. cFOS signals were counted on 5 serial sections of the designated spinal cord segment at a 120 microns interval between sections, and the average signal number was used as single data point for one mouse. We used ImageJ/FIJI for image contrast adjustment and signal quantification.

### *In vivo* distribution

C57BL/6J mice were administered with cy5-siRNA LNPs by intrathecal injection. The major organs including the heart, liver, spleen, lung, kidneys, brain, spinal cord, and lumbar L3-L5 DRGs were dissected from mice at different time points and imaged using an IVIS Lumina Series III Pre-clinical *in vivo* imaging system.

### Behavioral tests

A hot plate test was used to assess thermal sensitivity. The hot plate apparatus (BioSeb) was set to 52 °C. One mouse at a time was placed in the chamber and a stopwatch was started immediately. Time was recorded for hind paw licking, flinching, withdrawal response, or jumping. The test was terminated after 45 sec if the animal did not respond to avoid paw damage. For the capsaicin-induced acute pain model, an intraplanar injection of 0.01% capsaicin solution to the left paw was performed using a 31-gauge, sterile, single-use needle. The mice were then placed in a Palmreader behavior acquisition device to monitor spontaneous animal behaviors and paw surface contact recordings following the procedures defined in detail before^16^. To analyze nocifensive behaviors from our behavior recordings, we used ARBEL (Automated Recognition of Behavior Enhanced with Light), a supervised machine learning algorithm combining pose-estimation and light-based analysis of body part pressure and distance^63^. This approach enabled automated scoring of pain-related behaviors, including flinching and licking/biting, in freely moving mice. Behavioral classification was performed using ARBEL’s pre-trained licking/biting and flinching classifiers. For inflammation-induced pain, mice were subjected to 3% of Dextran Sulfate Sodium (MP Biomedicals 0216011080) into the drinking water for 4 days. Mice were habituated for at least two days before being placed into the recording device to monitor licking behavior. Licking behaviors were manually labeled frame-by-frame using a customized MATLAB-based scoring platform.

### *In vivo* toxicity evaluation

*In vivo* toxicity of TRPV1 siRNA LNPs was comprehensively studied. In brief, the major organs were harvested at the endpoint, sectioned, and H&E stained to evaluate the histological differences. In addition, blood was drawn, and serum was isolated at the end of the *in vivo* efficacy experiment. Various parameters, including ALT, AST and others, were measured to evaluate toxicity.

### Statistical analysis

A one-way or two-way analysis of variance (ANOVA) was performed when comparing three groups or more than three groups, respectively. Statistical analysis was carried out using Prism 10.0 (GraphPad). Data are expressed as standard deviation (S.D.) as described in the main text. All studies were performed at least in triplicate unless otherwise stated.

## Supplementary Materials

**Supplementary Figure 1.**
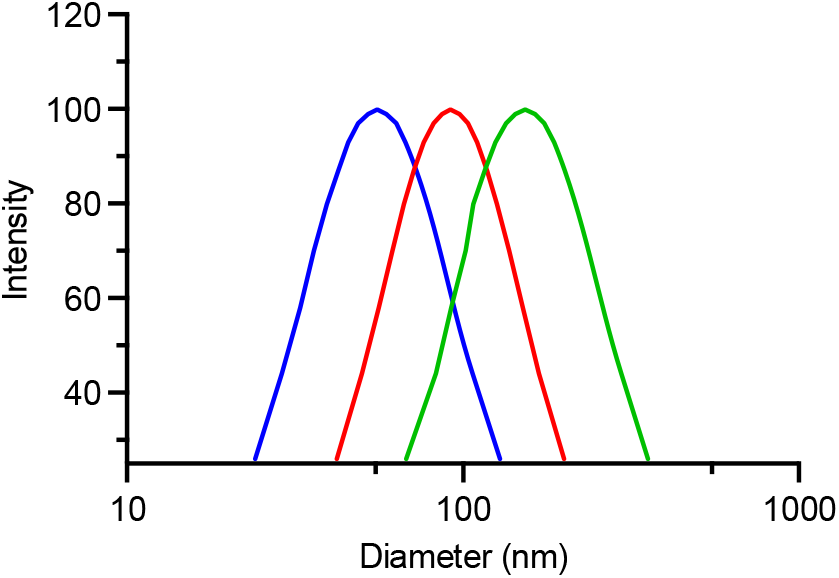
Particle sizes of various siRNA LNPs. The sizes were around 55 nm, 92 nm, and 153 nm, respectively, as determined using a ZETAPals analyzer.

**Supplementary Figure 2.**
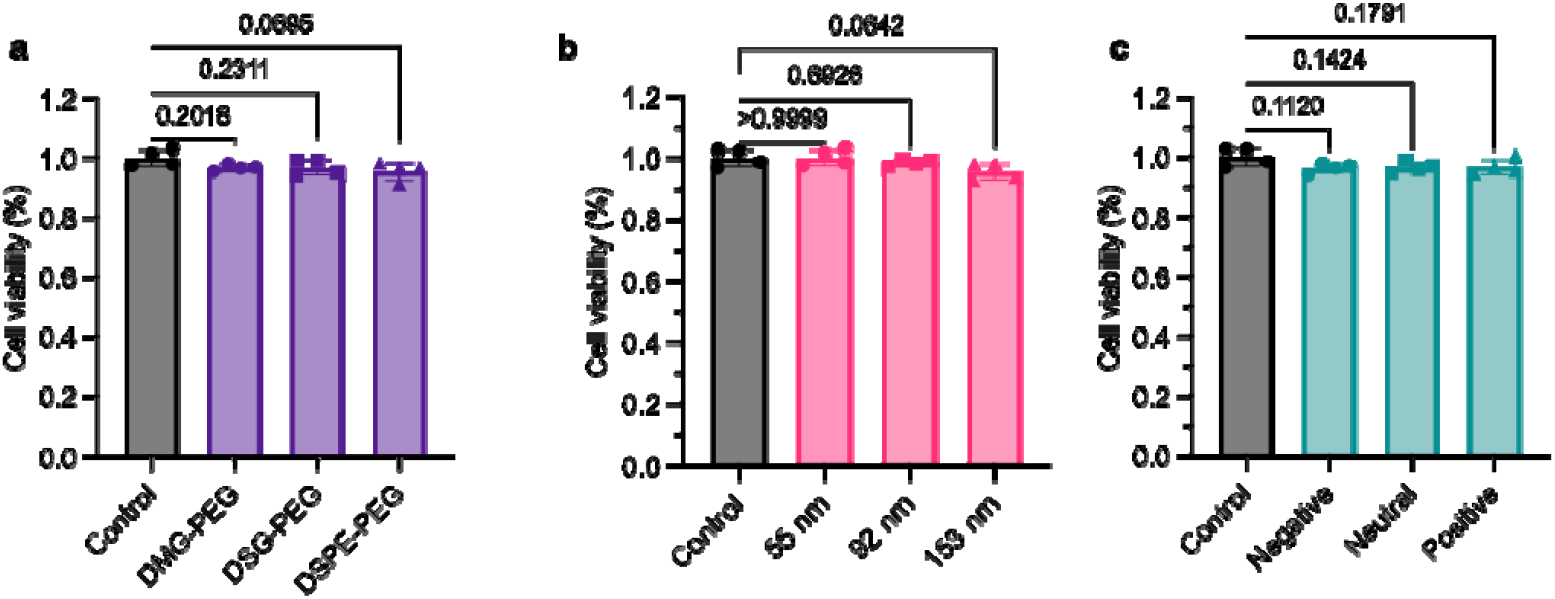
Cytotoxicity of siRNA LNPs. a-c, 175-luc cells were treated with LNPs with different lipid-PEGs (a), particle sizes (b), and charges (c), and the AlamarBlue assay was used to detect the cytotoxicity of these LNPs (N=4, mean ± SD, One-way ANOVA with Dunnett’s post hoc test).

**Supplementary Figure 3.**
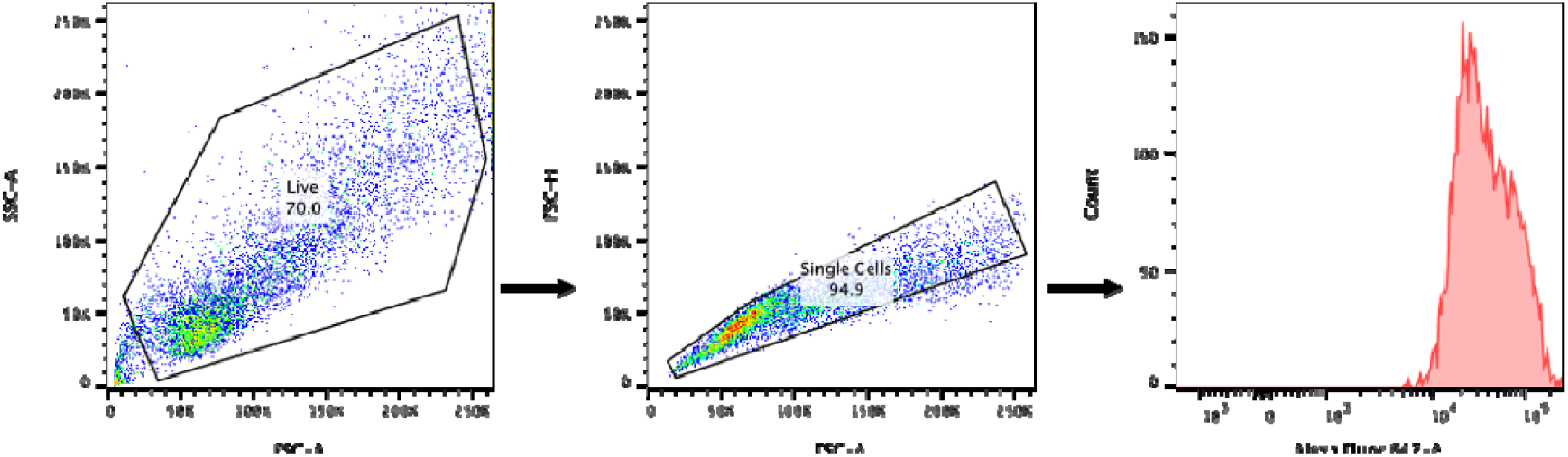
Flow cytometry gating strategy for cellular uptake of cy5-siRNA LNPs.

**Supplementary Figure 4.**
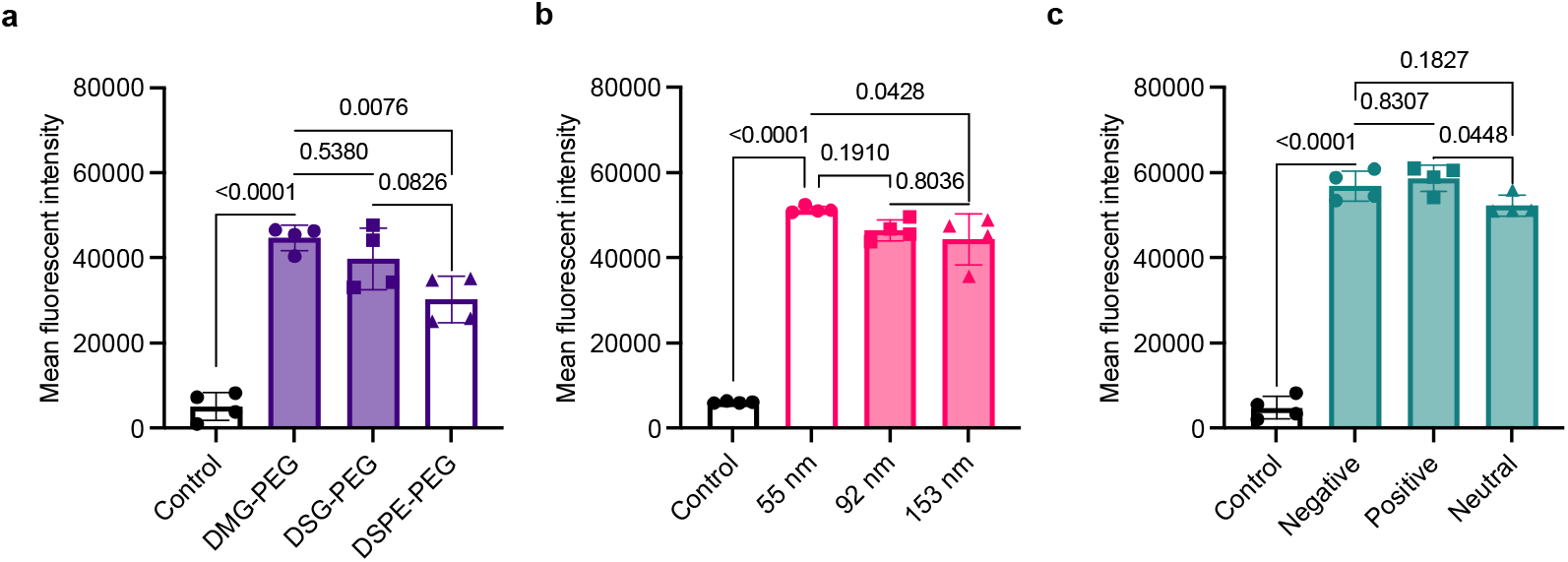
Flow cytometry quantification of the cellular uptake of cy5-siRNA LNPs. Mean fluorescent intensity of cellular uptake of cy-siRNA LNPs with different lipid-PEGs (a), particle sizes (b), and charges (c). 175-luc cells were treated with these cy5-siRNA-loaded LNPs for 24 hrs, and fluorescent intensity was measured by flow cytometry (N=4, mean ± SD, one-way ANOVA with Tukey’s post hoc test).

**Supplementary Figure 5.**
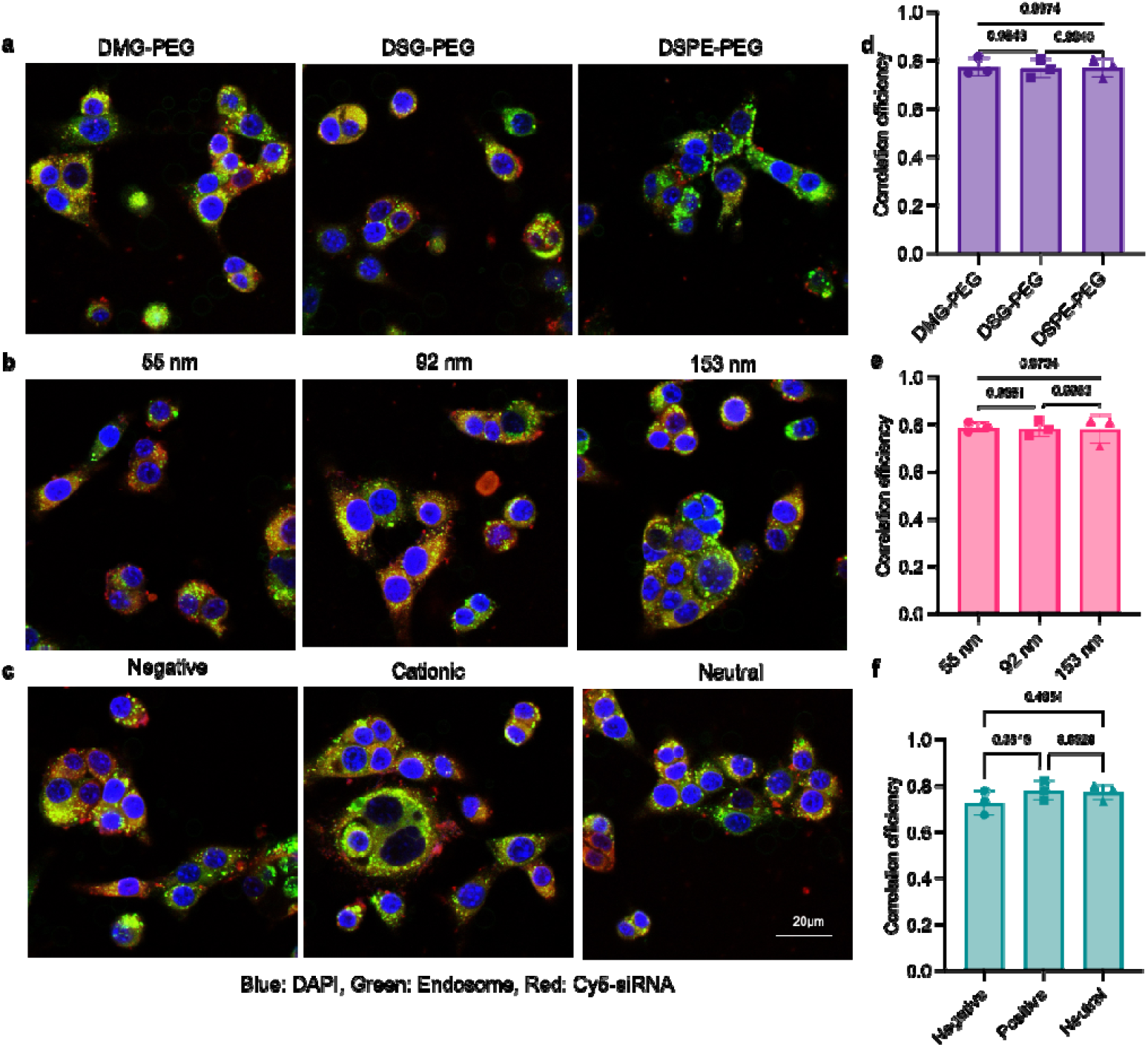
Endosome escape of different LNPs. Confocal images to show the endosome escape of different cy5-siRNA LNPs with different lipid-PEGs (a), particle sizes (b), and charges (c). (blue: DAPI, green: endosome, red: cy5-siRNA, scale bar =20 μm). 175-luc cells were treated with different cy5-siRNA LNPs for 24h, and then washed, fixed, and stained with Lysotracker and DAPI. Results showed that all the formulations mediated effective endosome escape. **d-f**, Quantitative analysis of the correlation efficiency of endosome (green) and cy5-siRNA (red) for **a-c** (N=3, mean ± SD, One-way ANOVA with Tukey’s post hoc test).

**Supplementary Figure 6.**
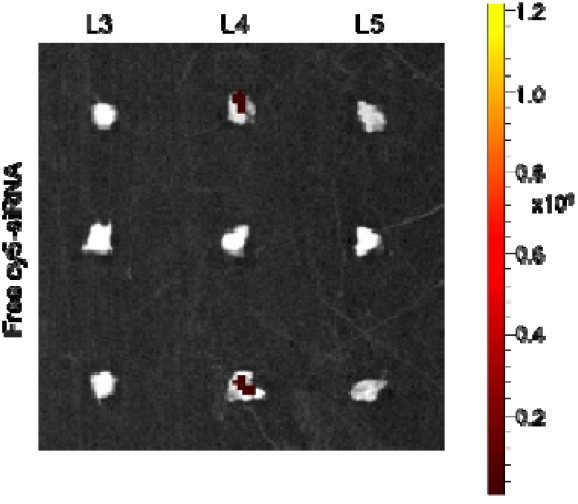
Free cy5-siRNA accumulation in the DRGs. C57BL/6J mice were administered with free cy5-siRNA via intrathecal injection, followed by L3-L5 DRGs imaging 24 h post-administration. The unit for the IVIS fluorescence intensity is (p/sec/cm^2^/sr)/(μW/cm^2^).

**Supplementary Figure 7.**
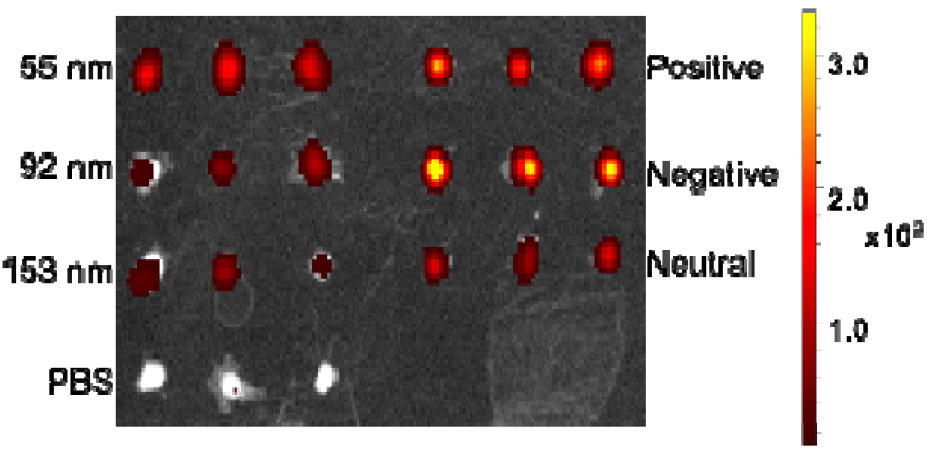
*Ex vivo* imaging of DRGs treated with cy5-siRNA LNPs with different sizes (left) and surface charge (right). The unit for the IVIS fluorescence intensity is (p/sec/cm^2^/sr)/(μW/cm^2^).

**Supplementary Figure 8.**
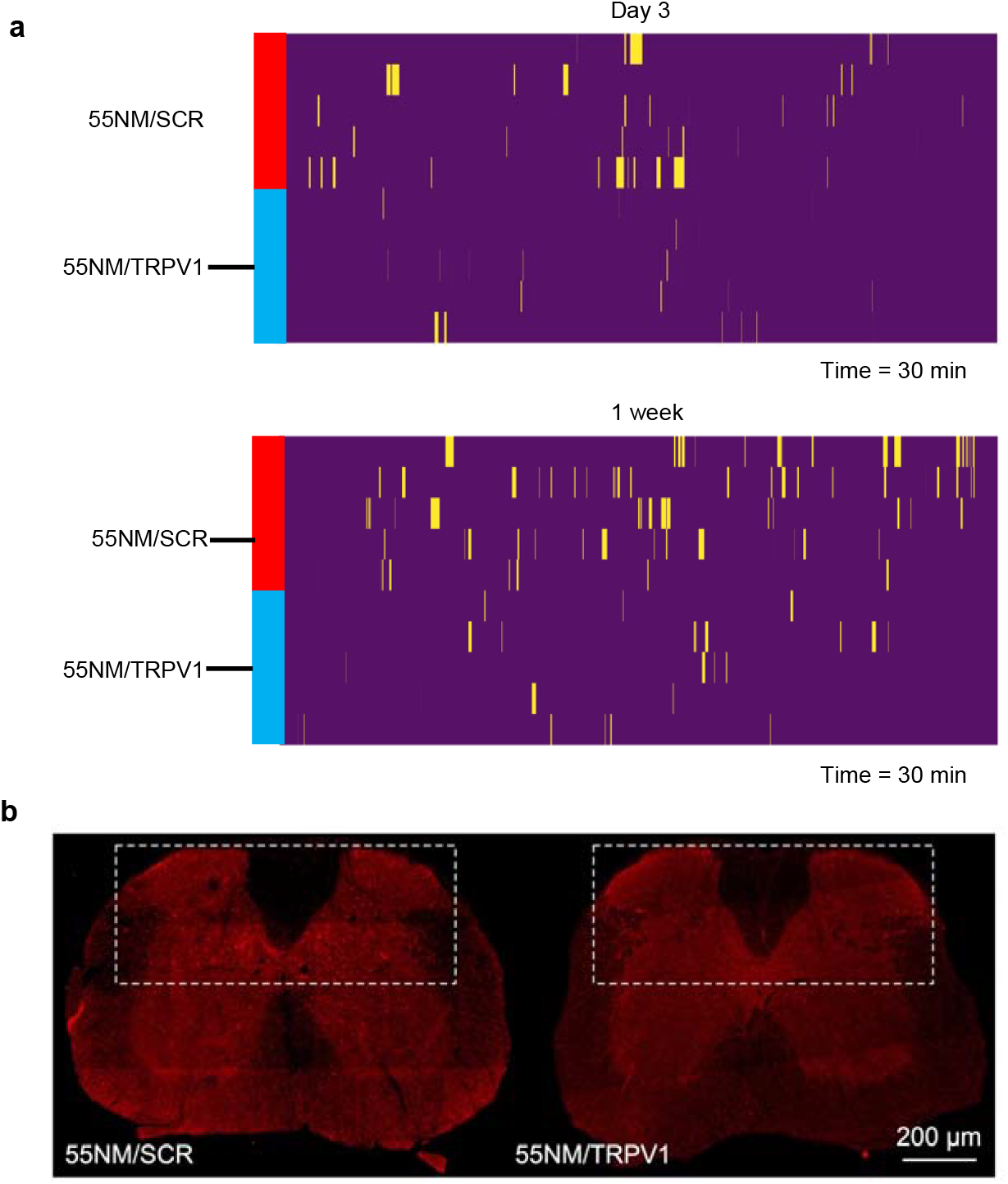
55NM/TRPV1 siRNA LNPs attenuated spontaneous licking bouts and spinal cord neural activation in mice driven by peripheral inflammation. **a**, Raster plots indicate individual licking bouts in both TRPV1 siRNA and scrambled LNPs treated groups at Day 3 and 1 week after intrathecal delivery. X-axis = 30 mins recording. N= 5 mice per group. **b**, Representative immunofluorescence images of spinal cord cross sections from mice with inflammatory colitis pain treated with 55NM/TRPV1 or 55NM/SCR. Scale bar = 200 μm. Dashed line indicates the area used for cFOS^+^ signal quantification.

**Supplementary Figure 9.**
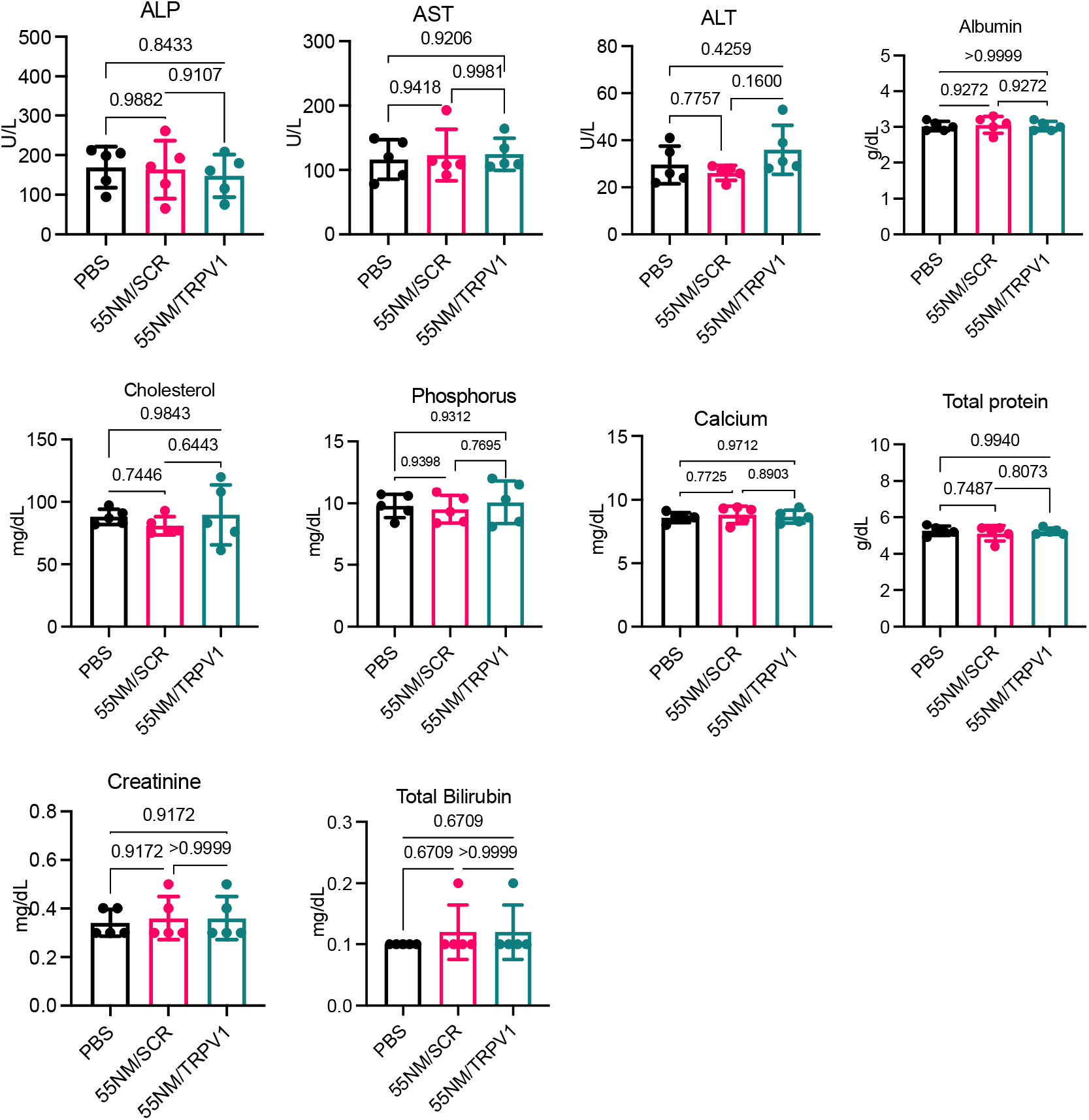
Blood chemistry analysis after various treatments. Blood serum was collected post-treatment with PBS, 55NM/SCR LNPs, and 55NM/TRPV1 LNPs (N=5, mean ± SD, One-way ANOVA with Tukey’s post hoc test).

**Supplementary Figure 10.**
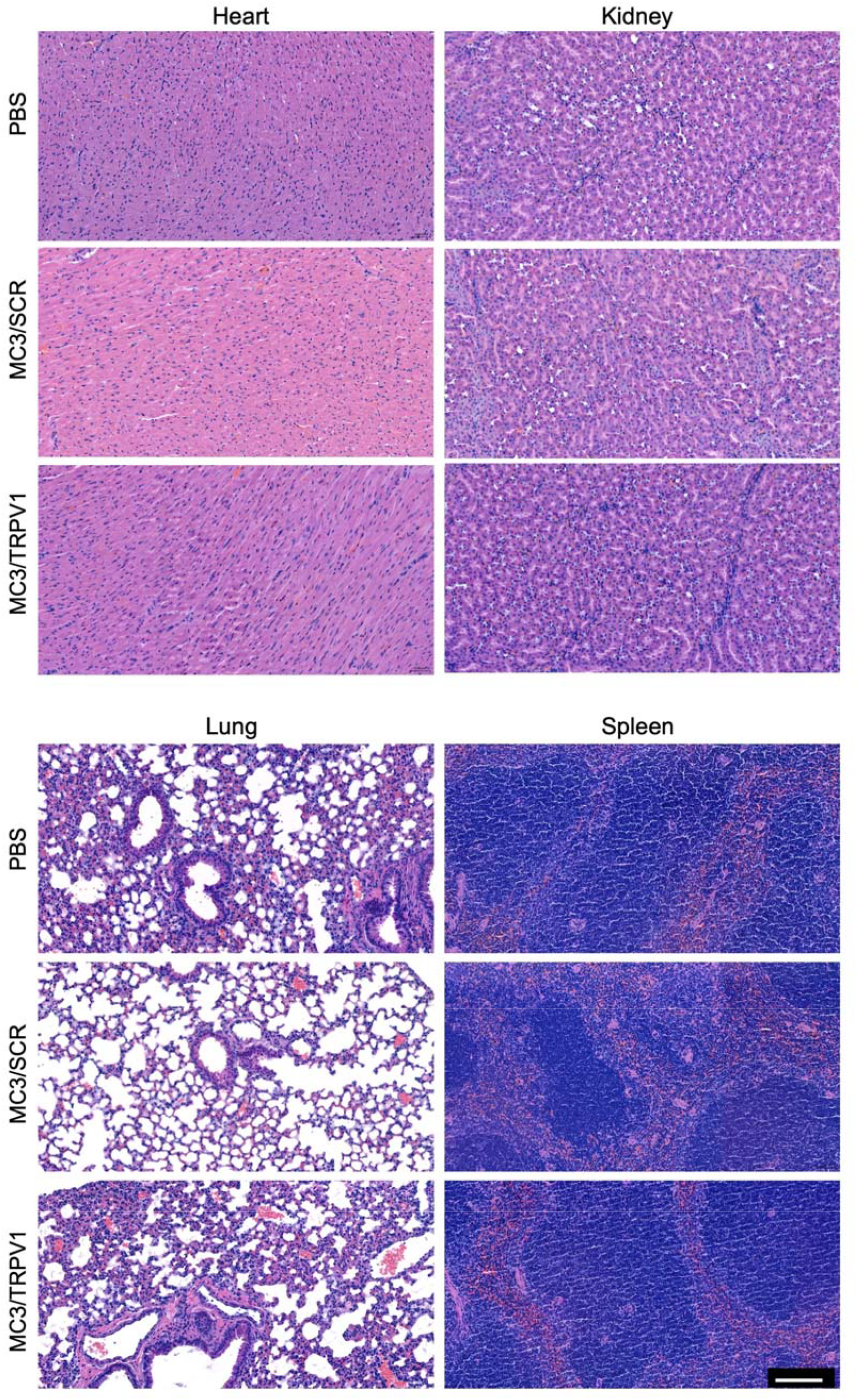
H&E staining images of the heart, kidneys, lung, and spleen. The organs were collected post treatment with PBS, 55NM/SCR LNPs, and 55NM/TRPV1 LNPs. Scale bar, 100 μm.

**Supplementary Table 1.**
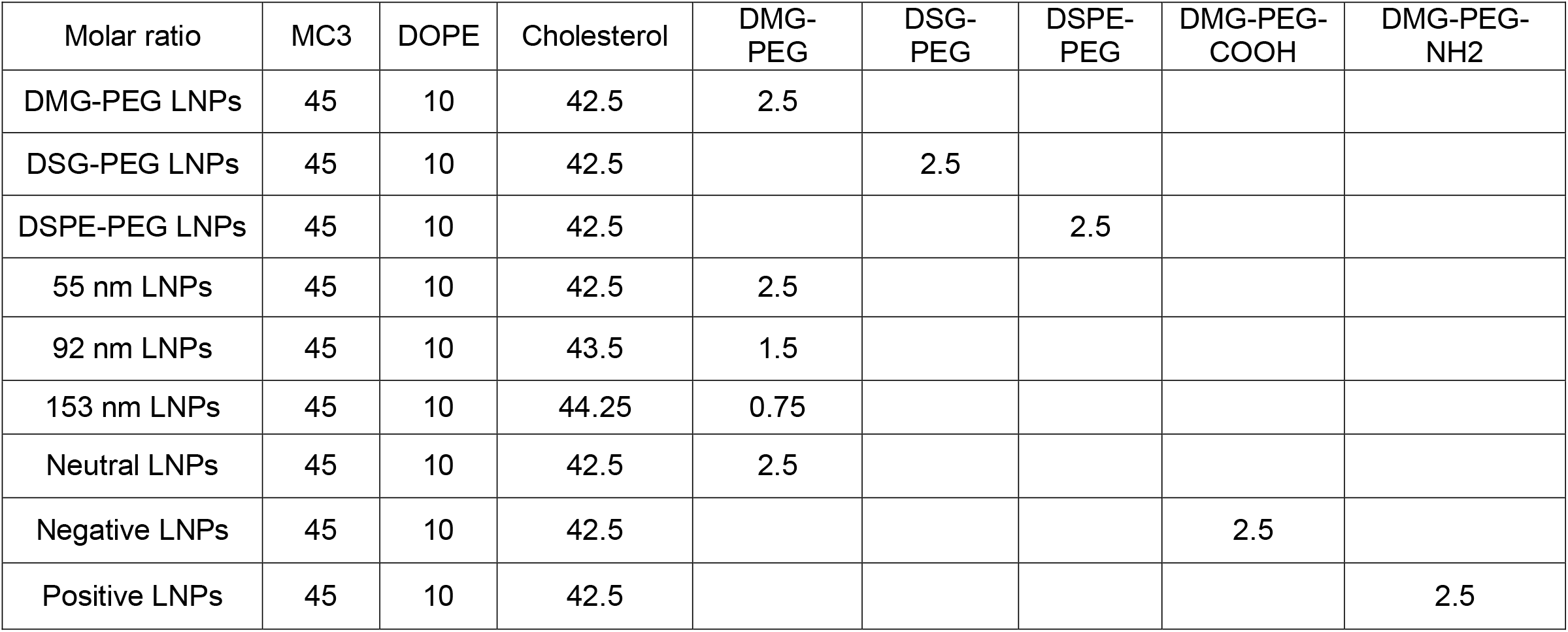
Lipid compositions for different siRNA LNP formulations.

## Acknowledgements

We thank Heena Aggarwal, Hae Lin Jang, and Yanhong Zhang for assistance with RT-qPCR. We thank the NeuroTechnology Studio at Brigham and Women’s Hospital for providing confocal microscope access and consultation on data acquisition and data analysis. This work was supported by the Basic/Clinical Collaborator Grant from the Department of Anesthesiology, Perioperative and Pain Medicine at Brigham and Women’s Hospital (PL and JS), and the National Institutes of Health grants R01HL159012 (JS), R01CA200900 (JS), R35NS105076 (CJW), and R01AT011447 (CJW). The content is solely the responsibility of the authors and does not necessarily represent the official views of the National Institutes of Health.

## Author Contributions

X.H., V.P., Y.S., contributed equally. J.S., C.J.W., P.L. directed the project and provided conceptual advice. X.H., V.P., Y.S. designed the research and performed the experiment and data analysis. X.H., Y.S., performed *in vitro* immunostaining and *in vivo* imaging experiments. V.P. designed, performed, analyzed and curated animal data and behavior describing noxious heat and capsaicin-induced mouse models. Y.C. performed the DSS inflammation pain experiment. X.J., H.Z., C.H., D.S.C., H.J.Y, A.J., W.R. helped with *in vitro* and/or animal experiments. B.Z. ran all animal videos through automated pipeline and curated data. O.B. created, tested and validated automated behavior algorithm. H.K. performed animal injections and behavior experiments. X.H., V.P., Y.S., P.L., C.J.W, J.S. drafted the manuscript. P.L., C.J.W., J.S. were involved in study conceptualization and supervision, manuscript writing and editing, and securing funding. All authors contributed to data analysis and writing of the paper. All authors have given approval to the final version of the manuscript. All the authors except CJW declare no competing financial interest.

